# Bidirectional Optogenetic Modulations of Peripheral Sensory Nerve Activity: Induction vs. Suppression through Channelrhodopsin and Halorhodopsin

**DOI:** 10.1101/2024.08.15.608055

**Authors:** Akito Kosugi, Ken-ichi Inoue, Masahiko Takada, Kazuhiko Seki

## Abstract

In this study, we investigated the potential of optogenetics for modulating activity of peripheral sensory nerves, particularly tactile and proprioceptive afferents, which are vital for movement control. Using adeno-associated virus serotype 9 vector, we selectively transduced channelrhodopsin (ChR2) and halorhodopsin (eNpHR3.0) into large-diameter sciatic nerve afferents of rats. Diverging from conventional dorsal root ganglion (DRG) approaches, we applied optical stimulation at the distal portion of the afferent nerve. The intensity of optical stimulation varied to modulate the extent of induction and suppression of afferent activity. Then, the effect of optical stimulation was determined by the activity recorded in the dorsal root of the same afferents. Our findings show successful induction and suppression of activity in large-diameter afferents via optical stimulation. By increasing the intensity of blue (for ChR2) and yellow (for eNpHR3.0) light stimulation, the activity of fast-conducting afferent fibers was preferentially evoked or inhibited in an intensity-dependent manner. These data indicate that the activity of large-diameter afferents can systematically be regulated by optogenetics. The present innovative methodology for manipulating specific sensory modalities at the nerve level offers a targeted and accessible alternative to DRG stimulation, expanding the therapeutic scope of optogenetics for treating sensory disorders.

## Introduction

Peripheral sensory nerve activity is integral in conveying sensory experiences to the central nervous system (CNS). Dysregulation of the peripheral sensory nerves, manifesting hyper-or hypoactivation, is a hallmark of various sensory disorders (1, 2). Conventional treatments for regulating peripheral nerve activity, such as pharmacological and electrical interventions (3−6), often lack specificity and may introduce adverse effects (4, 7, 8). The advent of optogenetics – a technique that manipulates neural activity with light-sensitive proteins – offers an alternative solution with unprecedented temporal and spatial precision that has already been established in the CNS (9−15).

In the present study, we attempted the extension of this optogenetic technique to peripheral sensory nerves, specifically targeting large-diameter afferents associated with tactile and proprioceptive sensations. The dorsal root ganglion (DRG) is the standard site for neural modulation (16−19), but is encumbered by its anatomical complexity and procedural risks (20, 21). By contrast, peripheral nerves are more accessible targets with reduced complexity, as evidenced by successful optogenetic manipulation of motor neuron axons (22− 25). The heterogeneity of peripheral sensory nerves, comprising afferents of varying modalities and diameters (26−29), introduces specific challenges for optogenetic application. Our previous work demonstrated the efficacy of adeno-associated virus serotype 9 (AAV9) vector for gene transduction into DRG neurons linked to fast-conducting primary afferent fibers (19, 30). In this study, we used the AAV9 vector to target selectively large-diameter sensory fibers in the sciatic nerve of rats transduced with a light-activated ion channel, channelrhodopsin (ChR2), and a light-driven chloride pump, halorhodopsin (eNpHR3.0). We found that optical stimulation of the sciatic nerve evoked or suppressed action potentials in its afferent fibers, with the magnitude of effect directly proportional to the light intensity. This suggests the feasibility of optogenetic approaches to fine regulation of sensory signals transmitted through peripheral nerve fibers.

## Results

### Transduction of ChR2 and eNpHR3.0 Using AAV9 Vector

We successfully transduced ChR2 (*n* = 7) or eNpHR3.0 (*n* = 10) into DRG neurons by intra-nerve injection of AAV9 vector (Fig. 1A and B). Using an acute experimental setup, we evaluated the effect of optical stimuli on the distal portion of the sciatic nerve (Fig. 1C and E).

**Figure 1.**
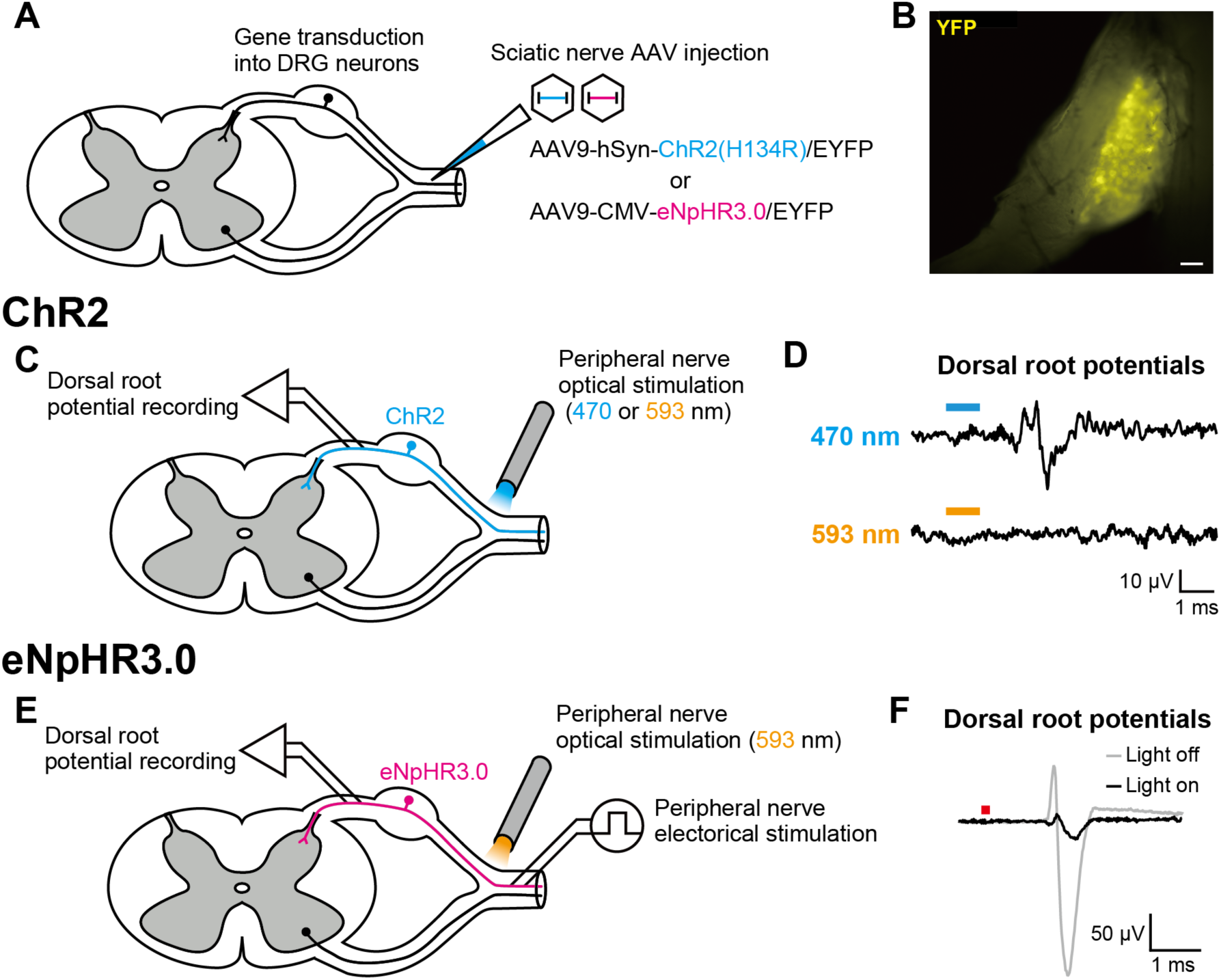
Optogenetic manipulation of sensory afferent activity by adeno-associated virus serotype 9 (AAV9)-mediated gene transduction into dorsal root ganglion (DRG) neurons and optical stimulation of nerve fibers. (A) Schema of AAV injection. Retrograde transport via sciatic nerve injection enabled gene transduction into DRG neurons. (B) Expression of YFP in the 5th lumbar DRG neurons. Scale bar: 200 μm. (C, D) Results on channelrhodopsin (ChR2) transduction. (C) Schema of the acute *in vivo* electrophysiological experiment. (D) Examples of dorsal root potentials evoked by light stimulation with different wavelengths. Waveforms represent the stimulus-triggered averaging of 20 responses. The bar above each waveform indicates the timing of light irradiation. (E, F) Results on halorhodopsin (eNpHR3.0) transduction. (E) Schema of the acute *in vivo* electrophysiological experiment. (F) Examples of electrically evoked volleys with (black line) and without (gray line) light irradiation. Waveforms represent the stimulus-triggered averaging of 20 responses. The red square above the waveforms indicates the timing of electrical stimulation.

### Optogenetic Induction and Suppression of Sensory Afferent Activity

In rats transduced with ChR2, the effect of blue light was examined by recording the conducting afferent volleys at the dorsal root (Fig. 1C). For rats transduced with eNpHR3.0, the nerve was pre-activated by electrical stimulation, and the effect of yellow light applied proximally to the stimulation site was then assessed at the dorsal root (Fig. 1E).

Representative results are shown in Figure 1D and F. Specifically, in the ChR2-transduced rats, 470-nm light produced long-lasting volleys in the dorsal root, while 593-nm light had no modulatory effect (Fig. 1D). In the eNpHR3.0-transduced rats, the volley size evoked by electrical stimulation (Fig. 1F, gray line) was significantly diminished by optical stimulation at 593 nm (Fig. 1F, black line). We found comparable induction or significant suppression of action potentials in all ChR2-(*n* = 7) and eNpHR3.0-transduced rats (*n* = 10; see the “Further Characterization of eNpHR3.0-Mediated Suppression” section for details). These findings demonstrated that light application could effectively evoke action potentials in sensory fibers of the ChR2-transduced rats and suppress sensory fiber activity in the eNpHR3.0-transduced rats.

### Modulations of Sensory Afferent Activity Dependent on Optical Stimulation Intensity

Next, we examined the impact of varying optical stimulation intensity on sensory afferent activity. In the ChR2-transduced rats, increasing the intensity of blue light from 20.5 to 51.6 mW resulted in a gradual increase in the amplitude of the recruited volley (Fig. 2A and B; significant main effect of light power analyzed by Friedman’s test, *χ*^2^(_6,60_) = 63.9, *p* < 0.0001). In eNpHR3.0-transduced rats, on the other hand, increasing the power of yellow light (≥ 8.7 mW) gradually decreased the amplitude of the electrically evoked volley (Fig. 2D and E; significant main effect of light power analyzed by Friedman’s test, *χ*^2^(_9,81_) = 73.8, *p* < 0.0001). These observations confirmed that optical stimulation could regulate the amplitude of recruited volleys by tuning the stimulation intensity. In addition, increasing the light power resulted in a shorter onset latency of the recruited volley in the ChR2-transduced rats (Fig. 2C) and a longer onset latency of the electrically evoked volley in the eNpHR3.0-transduced rats (Fig. 2F, latency suppression). The overall findings indicate that sensory afferent nerve activity can be finely controlled by adjusting optical stimulation parameters.

**Figure 2.**
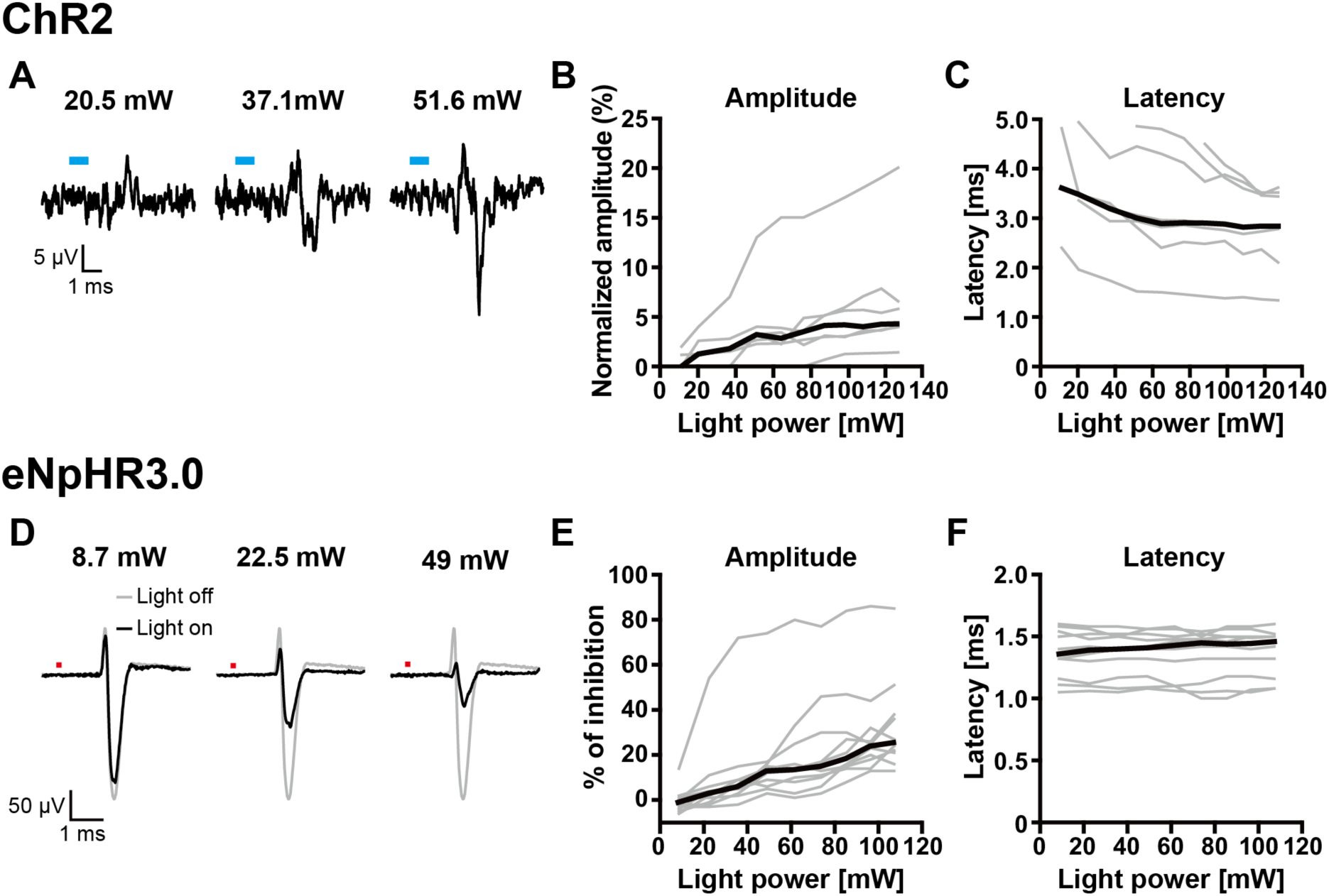
Effect of light power on optogenetic manipulation. (A−C) Results on channelrhodopsin (ChR2) transduction. (A) Examples of recruited volleys using different irradiated light powers. Waveforms represent the stimulus-triggered averaging of 20 responses. Blue bars above each waveform indicates the timing of light irradiation. (B, C) Changes in the amplitude and latency of recruited volleys as a function of light power. Each gray line represents one animal (*n* = 7), and thick black lines represent the median value of all animals. (D−F) Results on halorhodopsin (eNpHR3.0) transduction. (D) Examples of electrically evoked volleys using different irradiated light powers. Waveforms represent the stimulus-triggered averaging of 20 responses. Black lines indicate electrically evoked volleys with light irradiation and gray lines indicate volleys without irradiation. Red squares above each waveform indicates the timing of electrical stimulation. (E, F) Changes in the percentage of amplitude inhibition and latency of electrically evoked volleys as a function of light power. Each gray line represents one animal (*n* = 10), and thick black lines represent the median value of all animals.

### Selective Modulation of Large-diameter Afferent Activity

Next, we examined the afferent selectivity of optogenetic modulation, given the tropism of AAV9 vector for fast-conducting, large-diameter afferents (19, 30). In the ChR2-transduced rats, the lower power of optical stimulation exclusively recruited early-onset volleys, while its higher power additionally recruited late-onset volleys as well (Fig. 3A). Specifically, early-onset volleys were not observed at 11.4 mW irradiation, but became apparent at around 20.5 mW irradiation (Fig. 3B). By increasing the light power, late-onset volleys were first observed at around 64.7 mW (Fig. 3B). This outcome aligned with the expected large-diameter selectivity of AAV9 vector. We further compared the maximal amplitude between the early-onset (1.5–4 ms after stimulus onset; Fig.3A, red shading) and the late-onset (5–10 ms after stimulus onset; Fig.3A, blue shading) volleys. The earlier range corresponded with a conduction velocity of 15–45 m/s, which well fitted within the expected range for fast-conducting, large-diameter Aα/β fibers (> 14 m/s) (31). The results showed that the amplitude was larger in volleys with shorter latencies (Fig. 3C; Wilcoxon signed-rank test, *p* = 0.016). Therefore, we concluded that large-diameter afferents, potentially Aα/β fibers, were preferential targets for optogenetic induction and modulation in the ChR2-transduced rats.

**Figure 3.**
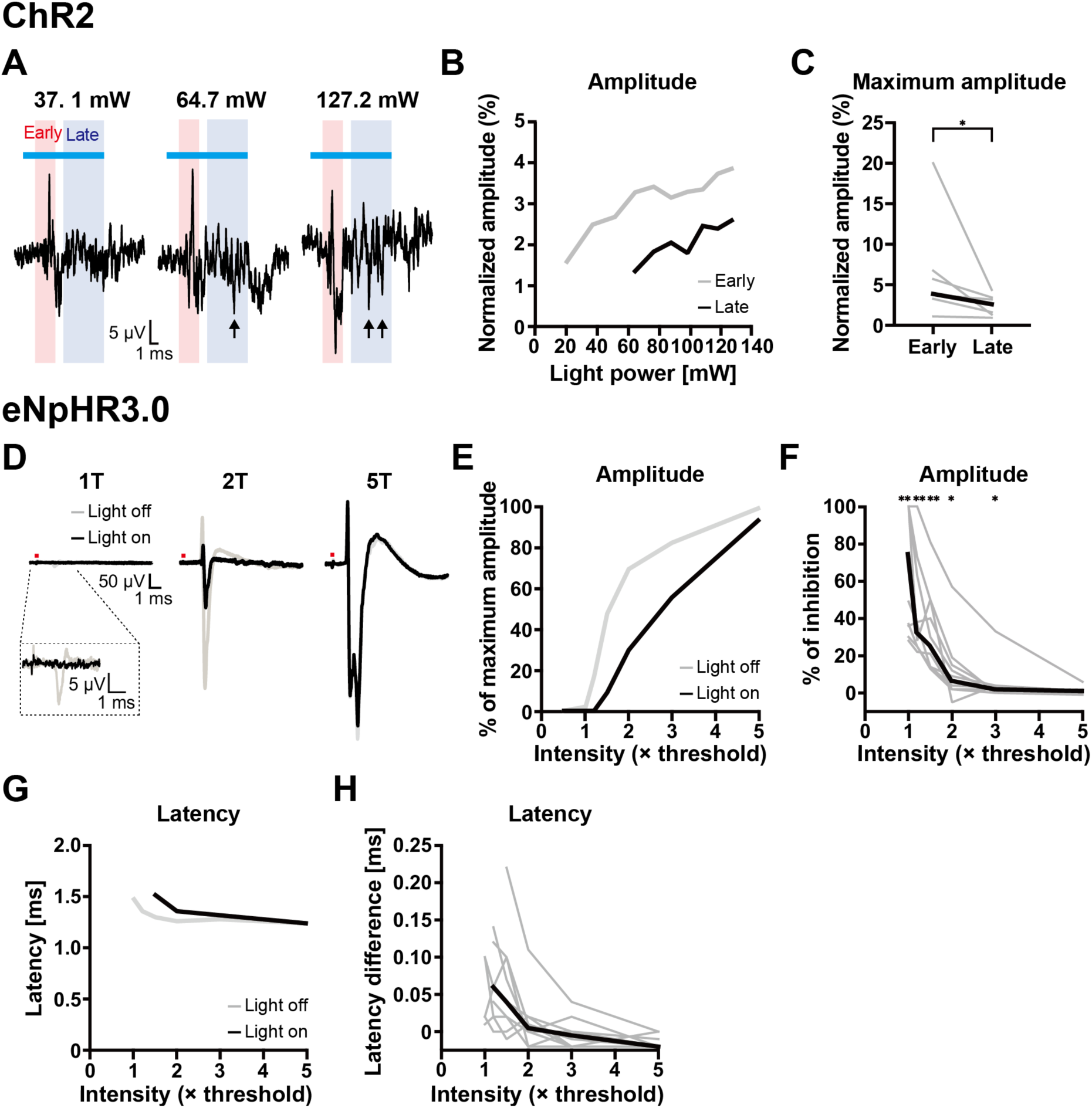
Large-diameter afferent preference. (A−C) Results of channelrhodopsin (ChR2) transduction. (A) Examples of recruited volleys using long light pulse durations. Waveforms represent the stimulus-triggered averaging of 20 responses. Red shading indicates the range of early-onset volleys (1.5−4 ms after stimulus onset), and blue shading indicates the range of late-onset volleys (5−10 ms after stimulus onset). Blue bars above the waveforms indicate the timing of light irradiation. Black arrows indicate individual peaks of late-onset volleys. (B) Changes in normalized amplitude of early- and late-onset volleys as a function of light power. The gray line indicates the early-onset component, and the black line indicates the late-onset component. Note that each line is plotted only at the light power at which the volley was observed. (C) Maximum normalized amplitude of early- and late-onset volleys evoked by long light pulse duration. Each gray line represents one animal (*n* = 7), and thick black lines represent the median value of all animals. \**p* < 0.05 (Wilcoxon signed-rank test). (D−H) Results of halorhodopsin (eNpHR3.0) transduction. (D) Examples of electrically evoked volleys using different intensities of electrical stimulation. Waveforms represent the stimulus-triggered averaging of 20 responses. Black lines indicate electrically evoked volleys using light irradiation and gray lines indicate volleys without irradiation. Red squares above each waveform indicates the timing of electrical stimulation. (E) Example of intensity–response curves of amplitudes with and without light irradiation from one animal. (F) Changes in the percentage of amplitude inhibition as a function of electrical stimulation intensity. Each gray line represents one animal (*n* = 10), and thick black lines represent the median value of all animals. \**p* < 0.05; \*\**p* < 0.01 (one-tailed Wilcoxon signed-rank test with Holm–Bonferroni correction compared to 0%). (G) Example of intensity–response curves of latency with and without light irradiation from one animal. (H) Difference in latency as a function of electrical stimulation intensity. Each gray line represents one animal (*n* = 10), and thick black lines represent the median value of all animals.

For the eNpHR3.0-transduced rats, we also confirmed preferential regulation of large-diameter afferents. We increased the intensity of electrical stimulation to determine how optical stimulation with the same parameters suppressed the size of electrically evoked volleys. We found a marked effect on volleys evoked by a stimulus current just above the recruitment threshold (1T in Fig. 3D). Notably, these lowest-threshold volleys were eliminated (100% amplitude inhibition) by optical stimulation in five of the 10 animals. By contrast, the optical stimulation did not eliminate the volley evoked at 2T, and the suppressive effect was less marked at 5T. We quantified the relationship between the suppressive effect and the electrical current in this animal (Fig. 3E). As reported earlier (32, 33), without optical stimulation, the volley size first increased rapidly and then slowly as a function of electrical current. With optical stimulation, however, this relationship changed from a non-linear to a more linear manner, and the size of the volley linearly increased as a function of current. This suggests the possibility that the suppressive effect may be afferent selective. For example, optogenetic suppression was more effective in the volley evoked at 2T (57%) than 5T (6%; ratio of gray line to black line in Fig. 3E). We calculated the efficacy of inhibition of the volley amplitude in all animals (Fig. 3F) and found that it was indeed larger for volleys evoked by a lower stimulus current. Maximal efficacy was obtained at around 1T and then decreased quickly, though it was still obvious until at 2T. The inhibitory effect was not statistically significant (Fig. 3F; one-tailed Wilcoxon signed-rank test, *p* = 0.063 without Holm–Bonferroni correction) despite that the stimulation intensity was increased to five times as much as the threshold level. We also found a comparable change in the onset latency of volleys. Indeed, the efficacy to generate a late-onset latency was decreased in volleys evoked by a higher stimulus current (Fig. 3G and H). These findings showed that optogenetic suppression had a greater effect on volleys evoked by lower-intensity electrical stimulation, suggesting preferential suppression of large-diameter afferent fibers. The overall results indicate that large-diameter sensory afferent activity can selectively be evoked and suppressed by optogenetics.

### Correlation Between Efficacy of Gene Expression and Extent of Optogenetic Modulation

We examined the relationship between the gene expression levels and the optogenetic modulation effects in both the ChR2- and the eNpHR3.0-transduced rats (Fig. 4). An example in Figure 4A shows recruited volleys and yellow fluorescent protein (YFP) expression in DRG neurons of two ChR2-transduced rats with varying expression levels. One animal exhibited larger efficacy in optical stimulation and stronger YFP signal intensity. We found a significant correlation coefficient between the maximal magnitude of recruited volleys and the YFP signal intensity (Fig. 4B; *r* = 0.99, *p* < 0.0001). A similar comparison was done in the eNpHR3.0-transduced rats (Fig. 4C). Again, a significant correlation coefficient was found between the amplitude inhibition percentage and the YFP signal intensity (Fig. 4D; *r* = 0.88, *p* = 0.001). It should be noted here that these significant correlations were maintained when an animal with notably high YFP expression was excluded (ChR2: *r* = 0.82, *p* = 0.047; eNpHR3.0: *r* = 0.68, *p* = 0.043). The results indicate the importance of consistent expression efficacy across animals for reproducible optogenetic modulation of sensory afferent activity.

**Figure 4.**
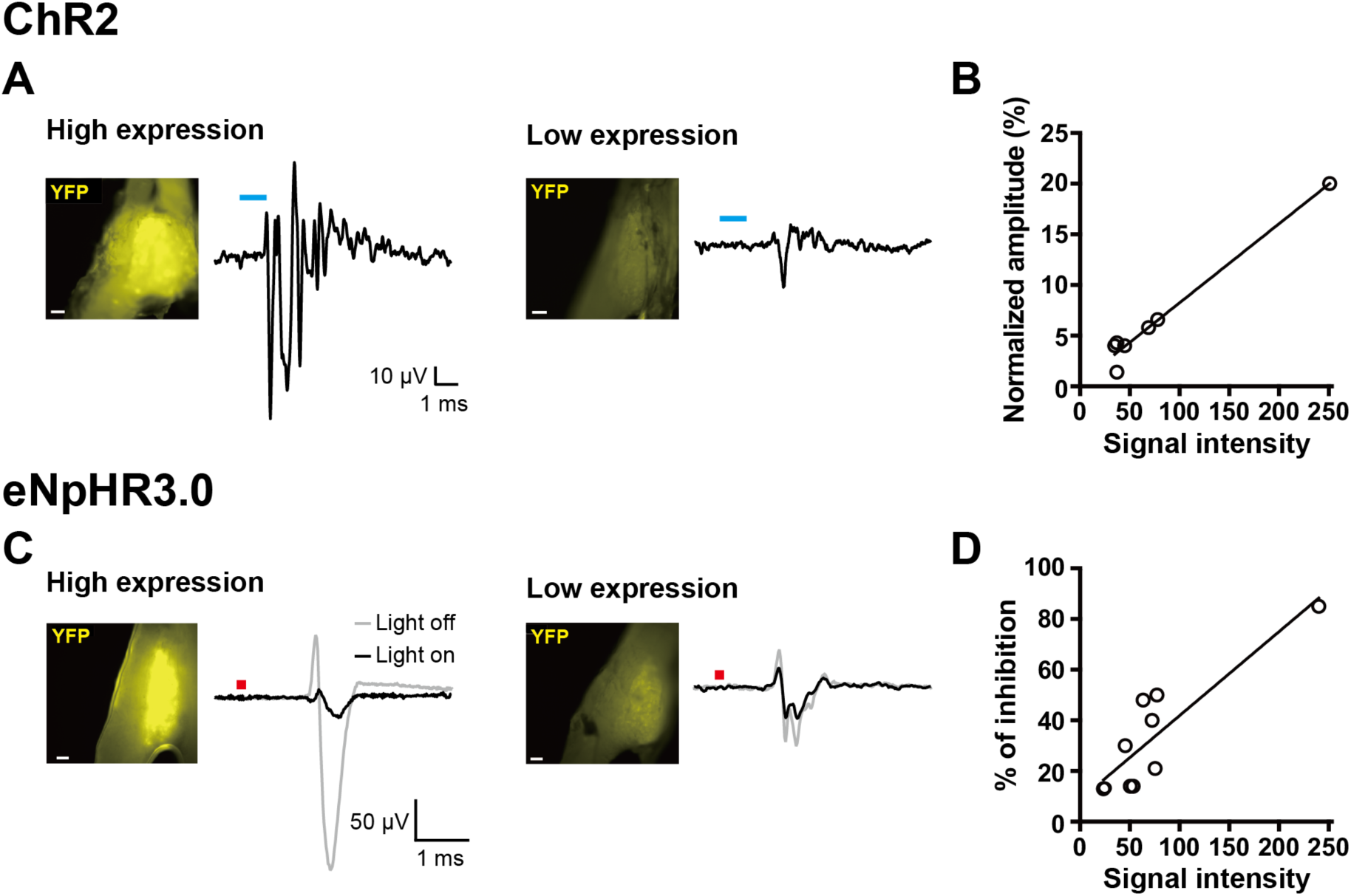
Correlation between optical efficacy and YFP intensity. (A, B) Results on channelrhodopsin (ChR2) transduction. (A) Examples of recruited volleys and YFP expression in dorsal root ganglion (DRG) neurons for two animals with different expression levels. Waveforms represent the stimulus-triggered averaging of 20 responses. Blue bars above each waveform indicates the timing of light irradiation. Scale bar: 200 μm. (B) Scatter plots and linear correlation between the normalized amplitude and YFP signal intensity of all animals (*n* = 7). (C, D) Results on halorhodopsin (eNpHR3.0) transduction. (C) Examples of electrically evoked volleys with (black line) and without (gray line) light irradiation and YFP expression in DRG neurons for two animals with different expression levels. Waveforms represent stimulus-triggered averaging of 20 responses. Red squares above the waveforms indicate the timing of electrical stimulation. Scale bar: 200 μm. (D) Scatter plots and linear correlation between the amplitude inhibition percentage and YFP signal intensity of all animals (*n* = 10).

Overall, our findings lead to similar conclusions in both of the ChR2- and the eNpHR3.0-transduced rats regarding the stimulation intensity-dependent modulation of large-diameter sensory afferent fibers and the correlation between the optogenetic effects and the gene expression efficacy. However, a notable difference was also observed between the two rat models, as described below (see the “DRG vs. Nerve: Comparative Analysis of Light Irradiation Sites” section).

### Further Characterization of eNpHR3.0-Mediated Suppression

Since this is the first study to inhibit action potential conduction at the peripheral nerve level, we further characterized the optogenetic suppression to determine a stable and reversible manner for inhibiting action potential conduction. The drop in the peak-to-peak amplitude of electrically evoked volleys occurred immediately after light irradiation (Fig. 5A, “Light”) and returned to baseline levels (“Baseline”) promptly after stopping light irradiation (Fig. 5A, “Recovery”). Significant differences were found between the “Baseline” and the “Light” session in a representative animal (Fig. 5B; one-way analysis of variance [ANOVA], *F*(_2,57_) = 4365, *p* < 0.0001; Tukey’s post hoc test, *p* < 0.0001), and all animals (Fig. 5C; Friedman’s test, *χ*^2^(_2,18_) = 15.2, *p* < 0.0001, Dunn’s post hoc test, *p* = 0.001). No difference was seen between the “Baseline” and the “Recovery” session (*p* = 0.61 in the representative animal and *p* = 0.99 in all animals), suggesting that light irradiation causes no side effects. In addition, there was no significant correlation between the elapsed time of light irradiation and the peak-to-peak amplitude in the “Light” session (Fig. 5A, *r* = −0.073, *p* = 0.76), indicating no time trend in this session. Furthermore, the onset latency of the volley in the “Light” session was significantly longer than that in the “Baseline” session in a representative animal (Fig. 5D; one-way ANOVA, *F*(_2,57_) = 742.3, *p* < 0.0001; Tukey’s post hoc test, *p* < 0.0001), and all animals (Fig. 5E; Friedman’s test, *χ*^2^(_2,18_) = 10.6, *p* = 0.002, Dunn’s post hoc test, *p* = 0.011). No difference was found between the “Baseline” and the “Recovery” session (*p* = 0.79 in the representative animal and *p* = 0.99 in all animals), suggesting the reversible nature of latency inhibition. These results indicate that optical stimulation can stably and reversibly inhibit action potential conduction of afferent fibers.

**Figure 5.**
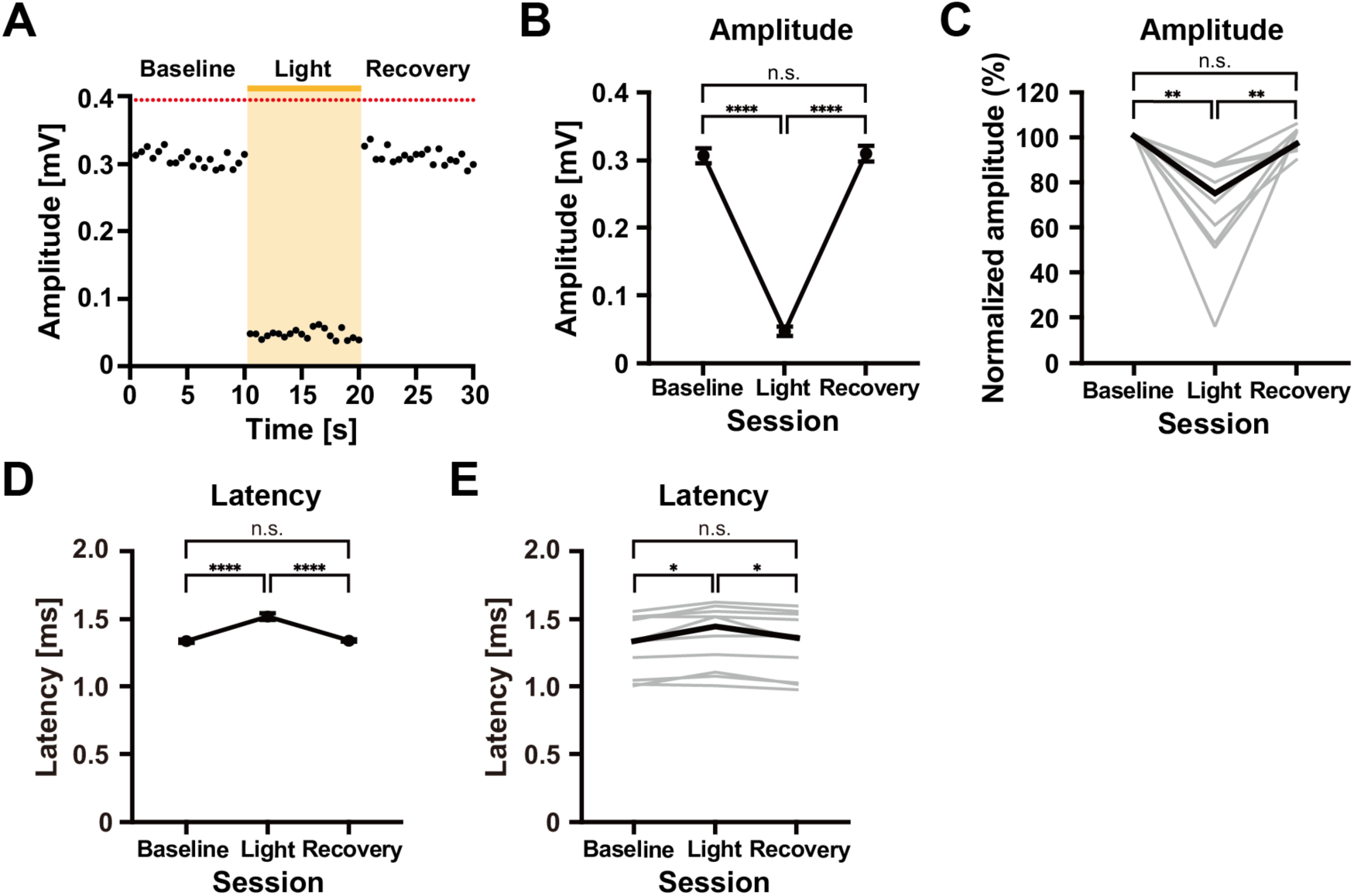
Characterization of optogenetic suppression. (A) Time course of peak-to-peak amplitudes of electrically evoked volleys. Red dots indicate the timing of electrical stimulation (2 Hz). The orange line and orange shaded area indicate the timing of light irradiation (10 s). (B, C) Effect of optical inhibition on the amplitude from a representative animal (B) and for all animals (C). For representative animal, data are represented mean ± standard deviation. For all animals, amplitudes in the “Light” and “Recovery” sessions were normalized by the average amplitude in the preceding “Baseline” session. Each gray line represents one animal (*n* = 10), and thick black lines represent the median value of all animals. (D, E) Effect of optical inhibition on latency from a representative animal (D) and for all animals (E). For representative animal, data are represented mean ± standard deviation. For all animals, each gray line represents one animal (*n* = 10), and thick black lines represent the median value of all animals. \**p* < 0.05; \*\**p* < 0.01; \*\*\*\**p* < 0.0001.

### DRG vs. Nerve: Comparative Analysis of Light Irradiation Sites

Building on previous studies that reported the optogenetic modulation of sensory afferent activity through light irradiation toward DRG neurons (7, 16−19), we next aimed to compare the efficacy of modulation between optical stimulation of DRG neurons and peripheral nerve fibers.

In the ChR2-transduced rats, we found that nerve-evoked volleys exhibited smaller amplitudes compared with DRG-evoked volleys (Fig. 6A–C). Since we applied the same intensity of optical stimulation in both conditions, this indicates that induction of sensory afferent activity by nerve irradiation is less effective than by classical DRG stimulation (Fig. 6C; Wilcoxon signed-rank test, *p* = 0.016).

**Figure 6.**
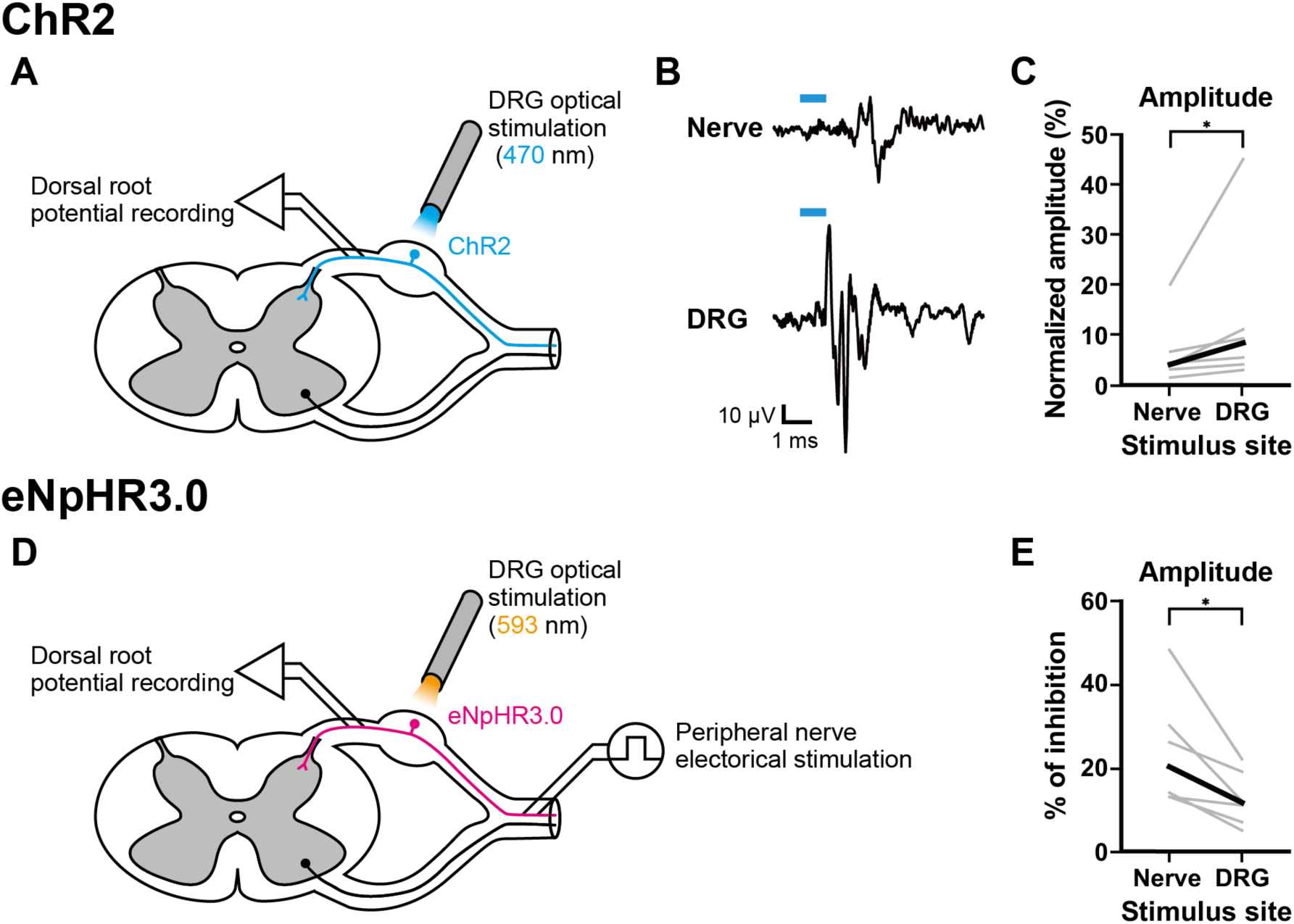
Sites and effects of optical stimulation. (A−C) Results on channelrhodopsin (ChR2) transduction. (A) Schema of the acute *in vivo* electrophysiological experiment. (B) Examples of recruited volleys by optical stimulation to the sciatic nerve and dorsal root ganglion (DRG). Waveforms represent the stimulus-triggered averaging of 20 responses. Blue bars above the waveforms indicate the timing of light irradiation. (C) The effect of the stimulus site on the amplitude. Each gray line represents one animal (*n* = 7), and thick black lines represent the median value of all animals. (D, E) Results on halorhodopsin (eNpHR3.0) transduction. (D) Schema of the acute *in vivo* electrophysiological experiment. (E) The effect of the stimulus site on the percentage of amplitude inhibition. Each gray line represents one animal (*n* = 6), and thick black lines represent the median value of all animals. \**p* < 0.05.

Contrastingly, in the eNpHR3.0-transduced rats, suppression of afferent fiber activity through nerve irradiation displayed the superior efficacy to DRG irradiation. This was demonstrated by more pronounced suppression of sensory afferent activity upon optical stimulation of the nerve compared with the DRG stimulation (Fig. 6D and E; Wilcoxon signed-rank test, *p* = 0.031).

These findings collectively suggest that while optical stimulation of sensory afferent fibers and DRG are capable of both inducing and suppressing their activity (as shown in Figs. 1–4), its effectiveness varies depending on the irradiation site. Specifically, in the eNpHR3.0-transduced rats, nerve irradiation exhibited more prominent suppression of activity compared with the classical DRG irradiation approach. Conversely, in the ChR2-transduced rats, the nerve application proved to be less effective in inducing sensory afferent activity. Such an efficacy difference highlights the significance of selecting the appropriate irradiation site to achieve the desired outcome in optogenetic modulation of sensory afferents. Notably, for induction of sensory afferent activity, the DRG may be a preferable choice, despite the increased complexity and surgical challenges associated with access thereto. This consideration is critical when weighing the trade-off between the efficacy and the procedural complexity in optogenetic manipulation of sensory pathways.

## Discussion

There are three points of novelty in the present study. First, we have succeeded in inducing and suppressing peripheral sensory nerve activity by optical stimulation. Second, the activity of fast-conducting, large-diameter afferent fibers was preferentially evoked or suppressed in a stimulation intensity-dependent manner. Third, the efficacy of optical stimulation varied depending on the irradiation site. In particular, the peripheral nerve was the preferred target for optogenetic suppression.

### Optogenetic Control of Peripheral Nerve Activity

In our study, we have found that optical stimulation can recruit primary afferent activity, with their amplitude and onset latency modulated in an intensity-dependent manner. A previous study reported that optical stimulation activated opsins expressed in axons at the node of Ranvier and generated action potentials (34). When the estimated conduction velocity of optically evoked volleys was investigated in the present study, it was consistently within the range of fast-conducting, large-diameter Aα/β fibers (> 14 m/s) (31). This implies the induction of Aα/β fiber activity by optical stimulation. In addition, our data confirmed recruitment of high-threshold late component volleys upon optical stimulation at high intensity with longer light-pulse durations, suggesting an off-target effect of optical stimulation (19). Nevertheless, the amplitude for late component volleys was usually smaller than for early component volleys (Fig. 3B and C). Thus, we can conclude that the optical stimulation in this study preferentially induces the activity of Aα/β fibers.

We also effectively demonstrated optogenetic suppression of axonal action potentials which were beforehand activated at the time of stimulation. When the action potentials were generated by electrical stimulation near the threshold current, we observed almost complete elimination of electrically evoked volleys. Electrical nerve stimulation is known to recruit action potentials in accordance with axonal diameter, as axons with larger diameters offer lower electrical resistance, making them more susceptible to depolarization in response to smaller input signals (35, 36). The earliest component of the recorded volleys, known as the “Aα” wave, reflects action potentials generated by group I muscle afferent fibers, which possess the largest axonal diameter among various categories of primary afferents (33, 37−39). Consequently, the complete elimination of volleys by optogenetic suppression in this study (Fig. 3D) indicates the superior effectiveness of optical stimulation in suppressing the activity of group I afferents. We also found the efficient optogenetic suppression of axonal action potentials primarily evoked by electrical stimulation at intensities twice as much as their recruitment threshold or below. It is well established that at such stimulation intensities, activities of both group I and II afferents are introduced, whereas finer afferents (like those of group III) are not activated (32). Therefore, we suggest that optogenetic manipulation in this study indeed inhibits the activity of large-diameter primary afferents, such as group I and II fibers.

Until recently, optogenetic suppression of primary afferent activity has exclusively been demonstrated by targeting small-diameter DRG neurons that are responsible for transmitting nociceptive signals (7, 17, 18, 40−42), with limited attention given to large-diameter neurons. In an effort to fill this gap, we previously attempted to transduce genes by intra-nerve injection of AAV9 vector (19, 30), known for its affinity toward medium-to-large-sized DRG neurons that are involved in conveying mechanoreceptive and proprioceptive signals (43). Here, we have further extended our approach using the AAV9 vector to achieve not only excitation (19), but also inhibition of large-diameter primary afferents including group I and II fibers.

### DRG vs. Nerve

Theoretically, there are four possible locations for light irradiation to activate opsins expressed in the peripheral sensory system: peripheral tissue, spinal cord, DRG, and peripheral nerve (5, 44). Among these, targeting the peripheral tissue has limitations. Light penetration is restricted to skin tissue when it is irradiated from the body surface (24), thereby confining the modulatory effect to skin receptors. The impact on deep primary afferent terminals, such as muscle spindles and Golgi tendon organs, is either impossible or incomplete. Similarly, targeting the spinal cord (from its surface) is inefficient due to the highly myelinated dorsal white matter which makes the light scattered (45, 46). Light intensity is likely to be significantly diminished before reaching the intermediate layer of the spinal cord where group I fibers terminate. In the present study, we targeted the remaining two candidates, DRG and peripheral nerve, and revealed that targeting the DRG yielded a superior efficacy for optogenetic induction of large-diameter primary sensory afferents, whereas targeting the peripheral nerve had the advantage of their suppression.

These observed differences in the effectiveness of optical stimulation between the DRG and the peripheral nerve suggest the involvement of distinct mechanisms. In the peripheral nerve, light irradiation activates opsins expressed in axons and changes the intra-axonal potential (34). By contrast, irradiation toward the DRG leads to activation of opsins expressed not only in axons, but also in somata. It was our expectation that the effect of light irradiation might be amplified by addition of the somal effect. This is because DRG neurons have active ion channels in their somata and can generate action potentials (47, 48), although these action potentials are generally believed to be rare. Also, hyperpolarization of the somata can influence the membrane potential at the T-junction of DRG, where axonal bifurcation occurs within DRG. Hyperpolarization at the T-junction may interfere with action potential propagation due to the possible impedance mismatch (49, 50). It seemed surprising that we obtained the expected result on optogenetic induction but the opposite result on optogenetic suppression, indicating a lower efficacy of DRG irradiation. Here, we propose two potential explanations for this outcome. First, the anticipated contribution of T-junction hyperpolarization may have been less influential than axonal hyperpolarization. Second, the effect of axonal hyperpolarization due to DRG irradiation may have been minor compared to nerve irradiation, stemming from structural differences between the two. Specifically, somata and satellite cells are located predominantly in the dorsomedial region of DRG, while passing fibers are more evenly distributed throughout the rest of the ganglion (51). Such a location of somata and satellite cells may result in scattering of the light when irradiated from the DRG surface, diminishing its intensity by the time it reaches the axons.

The superior efficiency of optical stimulation targeting the nerve, compared with targeting the DRG, is desirable for *in vivo* application of this technique. In the past, various chronic optical stimulation tools have been developed, such as nerve cuffs with LED or optical fibers (52−54), LED directly placed on the peripheral nerve (42), and even transdermal light irradiation to the peripheral nerve (24, 55). Conversely, methods for chronic optical DRG stimulation are limited *in vivo* (44) and are highly invasive, posing a risk of damaging DRG neurons during exposure at the stimulation site (20). Optical stimulation, especially for optogenetic suppression, at the peripheral nerve level opens new experimental possibilities of studying the involvement of sensory afferents in a variety of motor behaviors.

### Clinical implications

Damage to the CNS in the case of stroke and neurodegenerative diseases, including spinal muscular atrophy (SMA), lead to maladaptive changes in the periphery. In stroke, one common complication is spasticity, which results from hyperexcitability of the spinal reflex (56, 57). While it is assumed that reducing proprioceptive input to the spinal cord could help prevent the onset of spasticity, there has been no clinical treatment to date that offers selective and reversible manipulation of proprioceptive afferents. Similarly, SMA is characterized by motor neuron death and muscle atrophy, with reduced neuronal activity in spinal sensorimotor circuits as a secondary change (58, 59). Increasing proprioceptive input to the spinal cord may compensate for motor impairments in patients with SMA (60). In the present study, we have shown the feasibility of selective and reversible induction and suppression of large-diameter primary sensory afferent fiber activity using optogenetics. This novel technique has a significant potential for advancing clinical research works on and therapeutic approaches to conditioning spinal sensorimotor circuits, thereby offering new possibilities of targeted interventions against CNS damage caused by stroke and neurodegenerative disorders.

## Methods

### Ethical approval

All surgery was performed according to the institutional guidelines for animal experiments and the National Institutes of Health Guide for the Care and Use of Laboratory Animals. All experiments were approved by the experimental animal committee of the National Institute of Neuroscience (approval number 2022020).

### Experimental animals

Seventeen male Slc:Wistar rats (4 weeks old, 70–90 g) obtained from Japan SLC, Inc. (Shizuoka, Japan) were used. All animals were housed under a normal light/dark cycle (12 h:12 h), with the light on from 08:00 h until 20:00 h, in a temperature-controlled environment with food and water available ad libitum.

### Production of viral vectors

AAV9-hSyn-ChR2(H134R)/EYFP (2.5 × 10^13^ genome copies/mL) (19) and AAV9-CMV-eNpHR3.0/EYFP (1.8 × 10^13^ genome copies/mL) were produced using the helper-free triple transfection procedure and purified by cesium chloride gradient. Viral titer was determined by quantitative PCR using Taq-Man technology (Life Technologies, Gaithersburg, MD, USA). Vector purity was assessed by 4–12% sodium dodecyl sulfate–polyacrylamide gel electrophoresis and fluorescent staining (Oriole, Bio-Rad, Hercules, CA, USA). A transfer plasmid (pAAV-CMV-eNpHR3.0-EYFP-WPRE) was constructed by inserting the eNpHR3.0-EYFP fragment with a Woodchuck Hepatitis virus (WHV) posttranscriptional regulatory element (WPRE) sequence into an AAV backbone plasmid (pAAV-CMV, Stratagene, La Jolla, CA, USA).

### Vector injections

ChR2 (*n* = 7) or eNpHR3.0 (*n* = 10) were transduced into DRG neurons by retrograde transport following a sciatic nerve AAV injection (Fig. 1A). The protocol for AAV injection was based on previous studies (19, 30). Briefly, animals were anesthetized by intraperitoneal injection of medetomidine/midazolam/butorphanol anesthesia (0.2, 2.0, and 2.5 mg/kg, respectively). Adequate depth of anesthesia was monitored frequently by monitoring pupil size and flexion reflex to the paw pinch. After anesthesia, the animals were fixed in a prone position, and an incision was made in the left gluteal muscle. The sciatic nerve was identified below the biceps femoris muscle and isolated approximately 20 mm from the surrounding tissue. A 31-gauge needle tube was inserted 10 mm into the isolated nerve. The needle tube was connected by polyethylene tubing (JT-10, EICOM, Kyoto, Japan) to a Hamilton syringe (1702RN, GL Science, Tokyo, Japan) that was mounted into a microinjection pump (LEGATO 180, KD Scientific, Holliston, MA, USA). The tubing, syringe, and needle tube were filled with an electrically insulating stable fluorocarbon-based fluid (Fluorinert, 3M, St Paul, MN, USA). Then 1% Fast Green (0.6 μL) was added to the viral vector solution to visualize the injected solution. After insertion, the needle tube was left in place for 5 minutes to allow sealing of the tissue around the needle tip. After, 3 μL of viral vector solution was injected at a rate of 0.6 μL/min. The needle tube was removed at 10 minutes after the end of the injection to ensure absorption of the solution. Two separate injections were performed into the common peroneal and tibial branches of the sciatic nerve to ensure that the nerve was filled uniformly (in total, 6 μL of viral vector solution was injected). Finally, the wound was closed with a non-absorbable suture, and the animals were allowed to recover at 37°C.

### Acute electrophysiological experiments

To test whether ChR2 and eNpHR3.0 could be used to recruit and inhibit activities of large-diameter somatosensory afferents *in vivo*, an acute electrophysiological experiment was performed 4–8 weeks after AAV injection. First, animals were anesthetized by intraperitoneal injection of medetomidine/midazolam/butorphanol anesthesia (0.2, 2.0, and 2.5 mg/kg, respectively). After a tracheotomy, anesthesia was maintained by inhalation of 0.5–1.0% isoflurane throughout surgery. Electrocardiogram, oxygen saturation levels, and rectal temperature were monitored and maintained within a physiological range.

To expose the lumbar spinal cord, a laminectomy was performed from the 1st lumbar to the 1st sacral vertebrae. Then, the upper thoracic spine as well as the base of the tail were immobilized using clamps. The dura was opened with a sharp needle, and the 4th and 5th lumbar dorsal roots were separated from the other roots. For electrical and optical stimulation, the left sciatic nerve was exposed and isolated from the surrounding tissue. The skin and muscle around the exposed roots and nerves were raised and tied to form a pool, which was filled with mineral oil to protect the exposed roots and nerves from drying out. After surgery, anesthesia was maintained by thiopental sodium (10 mg/kg), and rocuronium bromide (10 mg/kg) was administered intravenously to achieve neuromuscular blockade.

To apply electrical stimulation, a bipolar cuff electrode was mounted on the isolated sciatic nerve. Stimulus current was generated using an isolator (SS 102L, Nihon Kohden, Tokyo, Japan) coupled to an electrical stimulator (SEN7203, Nihon Kohden) with various stimulus intensities (1–10-times threshold) at a fixed pulse width (100 μs). A multimode optical fiber (960 μm in diameter, 0.63 numerical aperture; Doric Lenses, Quebec, Canada) connected to a light source (LISER_LED470, Doric Lenses) was placed 10 mm proximal to the cuff electrode to ensure the stimulus site was not directly irradiated (Fig. 1E).

To record stimulus-evoked volleys, a bipolar hook electrode was mounted on the isolated dorsal root at 15–25 mm proximal to DRG. Recorded potentials were amplified (× 1000) and band-pass filtered (15–10 kHz) using an amplifier (MEG-6108M, Nihon Kohden), and digitized at 50 kHz using an analog/digital converter (Digidata 1550, Molecular Devices, LLC, San Jose, CA, USA). The average of 20 responses, consisting of a 10 ms pre-stimulus baseline period followed by a 100 ms post-stimulus period, was obtained. Then, the peak-to-peak amplitude and latency between onset of the stimulus and beginning of the response were calculated. The beginning of the response was defined as a time that exceeded the value of the mean ± 3× standard deviations of the background signal recorded 10 ms before the stimulation.

To confirm the effect of optical facilitation, optical stimulation of blue light at 470 nm was applied to the sciatic nerve. The stimulation was applied 20-times with a fixed stimulus duration (1 ms) at an interval of 0.5 s (2 Hz). To test whether the effect of optical stimulation was caused by activation of ChR2, irradiation at a different wavelength (593 nm, 1 ms duration) was also performed. To characterize the response properties of optical facilitation, intensity–response characteristics of the irradiated light power were examined by changing the intensity of light power (11.4–127.2 mW) at a fixed stimulus duration (1 ms). To test whether optical stimulation activated large-diameter afferents selectively, intensity– response characteristics with longer light pulse duration (10 ms) were further examined because longer and stronger optical stimulation can recruit high-threshold non-target fibers (19). The light power was measured at the tip of the optical fiber using a power meter (PM100D, Thorlabs, Newton, New Jersey, USA). To facilitate comparison across animals, volley amplitudes were normalized to the maximum amplitude of electrically evoked volleys (10-times threshold intensity).

To confirm the effect of optical inhibition, three experimental sessions were performed: a baseline session (“Baseline”, light-off condition), a session during irradiation of yellow light at 593 nm (“Light”, light-on condition), and a session immediately following light deactivation (“Recovery”, light-off condition). Each session consisted of 20 electrical stimuli at an interval of 0.5 s (2 Hz) and proceeded consequently without a break. Yellow light was continuously irradiated for 10 s in only the “Light” session. To facilitate comparison across animals, volley amplitudes in the “Light” and “Recovery” sessions were normalized against the averaged amplitude in the preceding “Baseline” session. Considering reported side effects of light irradiation, such as heating (14, 61−63) and light-off rebound effects (64), the reversibility of light irradiation was confirmed by comparing “Baseline” and “Recovery” sessions. Because prolonged light irradiation can decrease the efficacy of inhibition over time (63, 65, 66), the temporal stationarity of optical inhibition was confirmed by calculating the correlation between the elapsed time of light irradiation for each stimulus and the peak-to-peak amplitude of each volley in the “Light” session.

To characterize the response properties of optical inhibition, three sessions were repeated under various conditions. First, intensity–response characteristics of irradiated light power were examined by changing the light power intensity (8.7–107.3 mW) at a fixed electrical stimulation (1.5-times threshold intensity). Then, intensity–response characteristics of electrical stimulation were examined by changing the intensity of electrical stimulation (1– 5-times threshold) at a fixed light power (107.3 mW). The percentage of amplitude inhibition was calculated as the difference in normalized amplitude between “Baseline” and “Light” sessions. The latency difference was also calculated as the difference in latency between “Baseline” and “Light” sessions and compared in each condition.

At the end of the experiments, small electrolytic lesions were made in the dorsal root and sciatic nerve using a 40 μA direct current stimulation of the recording and stimulating bipolar hook electrodes for 40 s. Then, the animals were deeply anesthetized by intravenous injection of thiopental sodium (20 mg/kg) and perfused transcardially with phosphate-buffered saline (pH 7.4), followed by 300 mL of 10% neutral buffered formaldehyde. After perfusion, the lumbar region of the spinal cord together with the DRG and sciatic nerve were removed.

### Histological quantification

To check expression of ChR2 and eNpHR3.0, the fluorescent signal intensity of YFP was examined. Fluorescent images of entire DRG neurons were acquired using an inverted microscope (BZ-X710, Keyence, Osaka, Japan) using fixed detector gain values without sectioning. To calculate the fluorescent intensity, the images were converted to grayscale, and then mean gray values within the region of interest were determined manually.

### Statistical analyses

The Wilcoxon signed-rank test was performed for all two group comparisons. For group analysis, Friedman’s test was performed with light power or recording sessions (“Baseline”, “Light”, and “Recovery” sessions) as a within-subject factor. One-way ANOVA was performed with three recording sessions only for single animal analysis in the optical inhibition experiment. Post hoc analyses were performed using Tukey’s or Dunn’s tests for multiple comparisons. When testing intensity–response characteristics of electrical stimulation in the optical inhibition experiment, one-tailed Wilcoxon signed-rank test with Holm–Bonferroni correction was performed to compare the percentage of amplitude inhibition in each stimulus intensity to the hypothetical value (0% inhibition). For all correlation analyses, Pearson’s correlation coefficient was calculated. The level of significance was set at *α* = 0.05. All electrophysiological data were analyzed using MATLAB 2021b (MathWorks, Natick, MA, USA), all fluorescent images were analyzed using OpenCV 4.6.0 with python 3.8, and all statistical tests were performed using Prism 10 (GraphPad Software, San Diego, CA, USA).

## Acknowledgments

We thank Dr. Hidemasa Furue for their helpful comments. Dr. Wupuer Sidikejiang, Dr. Shinji Kubota, and Ms. Moeko Kudo for their technical assistance. This work was supported by a Grant-in-Aid from the Japan Society for the Promotion of Science (JSPS) (grant numbers JP19H05724, JP19H01092, JP26120003, and JP 23H05488 [to K.S.], JP22H04922 [to K. I.], and JP21K17633 [to A.K.]), and research grants from the Japan Agency for Medical Research and Development (AMED) (grant numbers JP21dm0207092, JP21dm0207066, and JP19ek0109216 [to K.S.], and JP21dm0207077 [to K.S. and M.T.]).

## Author contributions

A.K. and K.S. designed the study. K.I. and M.T. developed the viral vectors. A.K. performed the surgery and electrophysiological experiments. A.K. analyzed the data. A.K. and K.S. wrote a draft of the manuscript. All authors approved the final version of the manuscript.

